# The impact of interneurons on nonlinear synaptic integration in the neocortex

**DOI:** 10.1101/2020.06.12.149260

**Authors:** Christopher Dorsett, Benjamin Philpot, Spencer LaVere Smith, Ikuko T. Smith

## Abstract

Excitatory inputs arriving at the dendrites of a neuron can engage active mechanisms that nonlinearly amplify the depolarizing currents. Interneuron-mediated inhibition can modulate this active process, in a subtype-dependent manner. For example, dendrite-targeting inhibition is hypothesized to increase the amplitude of synaptic input required to activate voltage-dependent nonlinear synaptic integration. To examine how inhibition influences active synaptic integration, we optogenetically manipulated the activity of two different subtypes of interneurons: dendrite-targeting somatostatin-expressing (**SOM**) and perisomatic-targeting parvalbumin-expressing (**PV**) interneurons. In acute slices of mouse primary visual cortex, electrical stimulation evoked nonlinear synaptic integration that depended on *N*-methyl-D-aspartate (**NMDA**) receptors. Optogenetic activation of SOM neurons in conjunction with electrical stimulation resulted in predominantly divisive inhibitory gain control, reducing the magnitude of the nonlinear response without affecting its threshold. PV activation, on the other hand, had a minimal effect on dendritic nonlinearities, resulting in a small subtractive effect. Furthermore, we found that mutual inhibition among SOM interneurons was strong and more prevalent than previously thought, while mutual inhibition among PV interneurons was minimal. These results challenge previous models of inhibitory modulation of active synaptic integration. The major effect of SOM inhibition is not a shift in threshold for activation of nonlinear integration, but rather a decrease the amplitude of the nonlinear response.

## Introduction

The integration of excitatory synaptic inputs in neuronal dendrites involves passive properties and active (i.e. voltage-gated) mechanisms^1^. Active mechanisms have been implicated as an important contributor to synaptic integration in a number of behavioral contexts^2–6^, brain regions^4,7,8^, and animal species^9,10^. Furthermore, inhibitory interneurons are key components in sculpting and refining activity in cortical circuits^11–15^ and behavior^5,16–19^. Still, it remains unclear how inhibitory interneurons modulate active dendritic processes during synaptic integration.

Inhibitory neurons are diverse in their morphology and connectivity, suggesting that they may have correspondingly diverse roles in neural circuitry. Interneurons exhibit a wide variety of axonal projection patterns onto their pyramidal cell targets. For example, basket cells are known to predominantly target cell bodies^20^, while Martinotti cells target the apical dendritic tufts^21^, and chandelier cells target the axon initial segment^22^. Thus, inhibition can be either proximal or distal relative to the site of excitatory input, and this spatial relationship influences their interaction. Proximal inhibition is effective at diminishing the amount of charge propagated to the soma^23–25^. Distal inhibition has been shown to be less effective^25,26^ and could be overcome by larger excitatory inputs^24^.

Inhibition can influence active synaptic integration as well. The *N*-methyl-D-aspartate (**NMDA**) receptor is a key component in nonlinear synaptic integration in dendrites^27^. Computational modeling suggests that NMDA spikes are particularly sensitive to distal dendritic inhibition. When colocalized to the same dendritic segment, even small inhibitory conductances are capable of eliminating the nonlinear increase in membrane potential associated with NMDA spikes, while somatically placed inhibition had negligible effects on both the spike waveform at the dendrite and the excitatory post-synaptic potential (**EPSP**) magnitude experienced at the soma^28^. Several *in vitro* experiments have also reached the conclusion that, in the context of active dendritic integration, the effectiveness of distal inhibition is more potent than previously assumed^29–31^. Active dendritic mechanisms can be studied with whole-cell electrophysiological recordings due to a characteristic response to subthreshold (i.e. below action potential threshold) stimulation^29,30,32–35^. Stimulating dendrites with linearly increasing intensity results in an initial linear somatic response, until the point at which sufficient depolarization recruits active mechanisms on the dendrites and responses become supralinear due to the recruitment of active voltage-gated mechanism^32,36,37^.

These responses are known to be affected by inhibition in a location dependent fashion^29,31^. However, whether proximal or distal inhibition has a larger effect on dendritic non-linearities remains disputed. Additionally, it remains unclear how specific interneuron subtypes affect active dendritic synaptic integration though it is natural to expect interneuron effects to vary by their characteristic subcellular targeting. We manipulated two prevalent interneuron subtypes: somatostatin-expressing interneurons (**SOM**) and parvalbumin-expressing interneurons (**PV**). These two subtypes have distinct axonal projection patterns. Around 60% of PV interneuron synapses onto layer 2/3 pyramidal cells are found in the perisomatic and proximal dendritic regions^38^. In contrast, SOM interneurons are biased towards distal regions, sending more than 90% of their axonal projections to dendrites^21,38^. We used whole-cell patch clamp recordings, electrical stimulation of excitatory inputs to pyramidal neurons, and optogenetic activation of interneurons to characterize how distinct interneuron subtypes influence active dendritic integration.

## Methods

### Animals

All procedures involving animals were carried out in accordance with the guidelines and regulations of the US Department of Health and Human Services and approved by the Institutional Animal Care and Use Committee of the University of North Carolina. Transgenic mice that express the improved light-activated cation channelrhodopsin (hChR2/H134R, hereafter ChR2) and the fluorescent protein tdTomato (tdTom) in a cre-dependent fashion (Ai27, Jackson labs #012567), were crossed with animals expressing cre-recombinase under either the SOM promoter (Jackson labs #010708) or PV promoter (Jackson labs #017320). Resulting experimental animals thus have ChR2 and tdTom expression in either SOM or PV cells. Equal numbers of male and female littermates from each genotype were used for all experiments. Mice were housed in a temperature and humidity-controlled environment on a 12-hour light/dark cycle with ad libitum access to food and water.

### Slice preparation

Cortical brain slices were dissected from adult mice (ranging in age from post-natal day 30 to 76) of both sexes which were determined to be heterozygous for both ChR2/tdTOM (Ai27^+^) and either PV-cre or SOM-cre. Slices were generated as described previously^39^. Briefly, mice were anesthetized with pentobarbital sodium (40mg/kg) and, following the loss of corneal reflex and toe-pinch response, were transcardially perfused with chilled dissection buffer (containing, in mM: 87 NaCl, 2.5 KCl, 1.25 NaH2PO_4_, 26 NaHCO_3_, 75 sucrose, 10 dextrose, 1.3 ascorbic acid, 7 MgCl, and 0.5 CaCl) bubbled with 95% O_2_ and 5% CO_2_. Mice were decapitated, their brains rapidly removed, and 350 μm-thick coronal slices were cut in chilled dissection buffer using a vibrating microtome (VT1000S, Leica, Buffalo Grove, IL). Slices were quickly transferred to a holding chamber to recover at 35° C for 20 minutes in artificial cerebrospinal fluid (aCSF; containing, in mM: 124 NaCl, 3 KCl, 1.25 NaH2PO_4_, 26 NaHCO_3_, 1 MgCl, 2 CaCl, 1.25 ascorbic acid, and 20 dextrose) bubbled with 95% O_2_ and 5% CO_2_. Following 20 minutes recovery, the holding chamber was transferred to room temperature for a minimum of 40 minutes before slices were used. Recordings were made in a submersion chamber perfused with bubbled aCSF at 2 ml/minute with temperature maintained at 33° C. For some experiments, 50 μM picrotoxin (PTX) or 100 μm (2R)-amino-5-phosphonopentanoate (APV) was added to the aCSF.

### Electrophysiology

Patch clamp pipettes were pulled from borosilicate glass using a gravity-driven pipette puller (PC-10, Narishige, Tokyo, Japan). Pipette tip resistances ranged from 4.2 – 7.8 MΩ when filled with internal solution (containing, in mM: 135 K^+^ gluconate, 4 KCl, 10 HEPES, 10 Na_2_-phosphocreatine, 4 Mg-ATP, 0.3 Na-GTP, 0.025 Alexa Floura 594; pH adjusted to 7.25 with KOH, osmolarity adjusted to ~295 mmol kg^−1^ with sucrose as needed). Layer 2/3 neurons were visualized for whole-cell recording on an upright microscope (Axio Examiner, Zeiss, Thornwood, NY) using infrared differential interference contrast or by fluorescence-based targeting for Cre^+^ neurons. Neurons were recorded in current clamp configuration using a patch clamp amplifier (Multiclamp 700B, Molecular Devices, Sunnyvale, CA) and data was collected using pCLAMP 10 software (Molecular Devices). Following an initial pipette seal resistance of ≥ 1 GΩ, capacitive transients were minimized before manually breaking into the cell. Input resistance was monitored by test current pulses. Cells were discarded if series resistance was initially > 25 MΩ or if either series or input resistance change by > 25% throughout the duration of recording. The bridge was rebalanced as necessary. Layer 2/3 pyramidal cell identity was confirmed by analysis of intrinsic membrane properties, responses to optogenetic stimulation, firing patterns to depolarizing current steps, and/or the presence of dendritic spines and apical dendrites after being filled with Alexa Fluor 594. Interneuron subtypes were identified by fluorescence, intrinsic membrane properties, response to optogenetic stimuli and firing response to depolarizing current steps.

For dendrite-dependent non-linearity experiments in layer 2/3 pyramidal cells, synaptic stimulation was performed as follows. After achieving a whole-cell recording configuration, the fluorescent signal from the Alexa Floura 594 was used as a guide to visually place a borosilicate theta stimulating pipette (World Precision Instruments, Sarasota, FL) filled with aCSF in close proximity (~5 μm) to the dendritic arbor of the cell being recorded from. Alternatively, if the dendritic arbor could not be visualized, the stimulating pipette was placed ~125 μm away from the soma (**Figure 1A** for histogram). Afferent axons from nearby cells could then be electrically stimulated (0.1 ms duration of negative current at various stimulus intensities, repeated for 5 sweeps) to elicit dendritic spikes. The stimulus intensity (SI) value required to produce a somatically detectable excitatory post-synaptic potential (EPSP) response was cell dependent, and ranged from 20μA to 240 μA with a median of 40 μA (mean = 46.363 ± 4.731 μA, n = 55). Once a detectable (i.e. ~0.5 mV) EPSP was achieved, the SI value was linearly increased by 10 or 20 μA steps until one of three scenarios was achieved: a clearly non-linear increase in EPSP value occurred, after which at least three additional SI values were recorded for curve-fitting analysis; the cell began to fire action potentials; or a depolarization of >35 mV occurred. The number of stimulus intensity values used to achieve these criteria range from 8 to 20 with a median of 11. To test for potential confounding effect of linearly increasing the SI value, in a subset of cells, SI values were presented in decreasing intervals. No differences in EPSP values were present between these trials and trials in which the intensity was linearly increased (**Supplemental Figure 1**). To evaluate the effect of optogenetic activation of a given interneuron subtype on dendrite-dependent non-linear increases in EPSP values, a 100 ms pulse of 450 nm light was delivered across the surface of the slice via a reflected laser pulse (peak power: 2.5 W; range: 2.21 – 6.25 mW/mm^2^; Techood Laser, Shenzhen, China). When electrical stimulation of nearby axons was paired with optogenetic stimulation, the electrical pulse was initiated 50 ms after the onset of a 100 ms light pulse. For these experiments, each SI value was repeated twice per sweep, once under control conditions and then again at the mid-point of the 100 ms light pulse.

**Figure 1.**
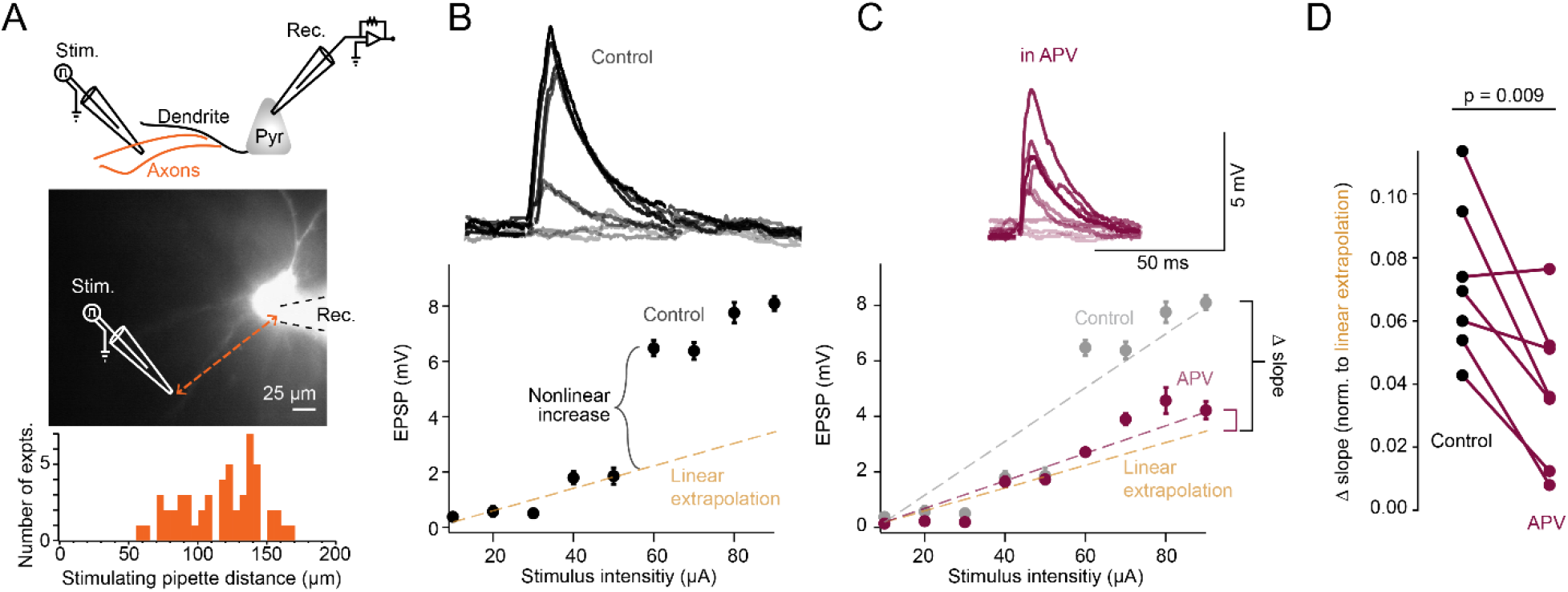
Electrical stimulation of afferent axons results in NMDAR-dependent dendritic supralinearities. (A) Top, cartoon schematic of recording configuration. Middle, example infrared image of layer 2/3 pyramidal cell filled with fluorescent Alexa 594 dye. Recording patch pipette outlined for illustrative purposes. Approximate location of theta glass stimulating pipette indicated. Bottom, Histogram of distance of each stimulating pipette from the recorded cell soma. (B) Example input-output (IO) curve showing subthreshold excitatory response to linearly increasing stimulus pulses (100 μs duration). Dashed line indicates linear extrapolation of mean EPSP values before responses become supralinear. Inset, example voltage trace responses. Error bars indicate ± s.e.m. (C) Same as (B) in the presence of 100 μM APV. Color-coded dashed lines indicate linear fit of entire IO function. (D) Change in slope (mV/μA) for entire IO function in control aCSF and in the presence of APV (n = 8).

### Immunofluorescence (IF)

Animals were anaesthetized with a mixture of ketamine (100mg/kg) and xylazine (15 mg/kg) and intracardially perfused with phospho-buffered saline (PBS) followed by 4% paraformaldehyde. After fixing overnight, 50 μm sections were cut and rocked in a blocking buffer (containing: 0.02% Sodium Azide, 0.03% Bovine Serum Albumin, 0.05% Goat Serum, 0.2% Triton X-100; in 250 mL PBS) for 1 hour. Primary antibody solutions were prepared in PBS using rabbit anti-RFP (Rockland 600-401-379; 1:400) and rat anti-SOM (Millipore Sigma MAB345, 1:400) primary antibody solutions were added to slices and incubated overnight at 4° C. Sections were then washed in blocking buffer at room temperature 3 x for 15 minutes each after which secondary antibodies were prepared using goat anti-rabbit (Invitrogen A10520; 1:500) and goat anti-rat (Invitrogen A11006; 1:500) were washed onto sections for 2 hours at room temperature. Sections were then washed in blocking buffer at room temperature 2 x for 15 minutes then once more in PBS containing 0.1% Tween20 for 15 minutes. DAPI staining (1:1000 dilution in PBS) occurred at room temperature for 15 minutes followed by a final wash in PBS at room temperature for 15 minutes. Sections were then mounted and imaged.

### Analysis

Data was analyzed using custom scripts for IGOR Pro. analysis software (WaveMetrics, Portland, OR), including event detection and analysis routines written by T. Ishikawa (Jikei University). To ascertain if electrically induced non-linear responses were dependent on NMDAR activation, cells IO plots were fit to the linear (*y = a + bx)* in control aCSF and aCSF containing 100 μm APV and the slope of the linear fit between the two experimental conditions was compared. One cell exhibited a sublinear IO curve in both control and APV containing aCSF and was excluded from further analysis (**Supplemental Figure 2**). Cells were analyzed for non-linearity by comparing mean somatic EPSP responses to a linear extrapolation of previous mean values to determine the non-linearity relative to linear extrapolation (NRLE) ratio^30^. Briefly, input-output (IO) plots of mean EPSP vs SI values were generated, and the SI values with the largest difference in EPSP responses (i.e. largest Δ) were identified. All prior EPSP response means were then fit to the linear. The experimentally derived EPSP for the SI value corresponding to the largest Δ was then compared to the expected value based on the linear extrapolation. Cells that had at least one experimental EPSP value that was at least 134% above the expected value were considered to display a non-linear response profile, while cells that did not were considered linear and were excluded from analysis (a total of 14/69 cell were linear – example cells in **Supplemental Figure 3**). If a cell displayed multiple points of non-linearity, the first instance was considered for analysis (example cells with multiple non-linear events in **Supplemental Figure 4**). In order to access changes in the magnitude of the dendritic dependent non-linearity caused by SOM and PV interneuron activation, the difference between the experimental EPSP values and the linear extrapolation at the SI with the largest Δ were compared under control conditions and during optogenetic activation. Furthermore, the average of the first three supralinear EPSP response values were compared under control stimulation conditions and during optogenetic activation of each interneuron subtype. In order to access changes in gain and offset, the entire IO curves under control and optogenetic conditions were fit to a sigmoid: base + {*max*/(1 + *exp*((*xhalf* − *x*)/rate)} where base and max are the baseline and maximal responses, respectively, and rate determines the slope parameter^31^. From this fitting we were able to calculate the degree of separation along the x-axis between control and optogenetic conditions from the x-half parameter. Furthermore, changes in slope due to optogenetic activation could be accessed by comparing the peak of the first derivative of the sigmoidal fit during control and optogenetic conditions.

### Statistics

Unless otherwise stated, all measurements are presented as mean ± s.e.m. Randomization and experimental blindness were not used for electrophysiology data as each cell serves as an internal control (e.g. EPSP value during control stimulation or in the presence of optogenetic activation of interneuron subtypes, spiking fidelity in standard aCSF or aCSF containing [50 μm] PTX, etc.). Statistical differences between control conditions and during optogenetic activation of interneuron subtypes were accessed by repeated-measures t-tests with α = 0.05

**Figure 2.**
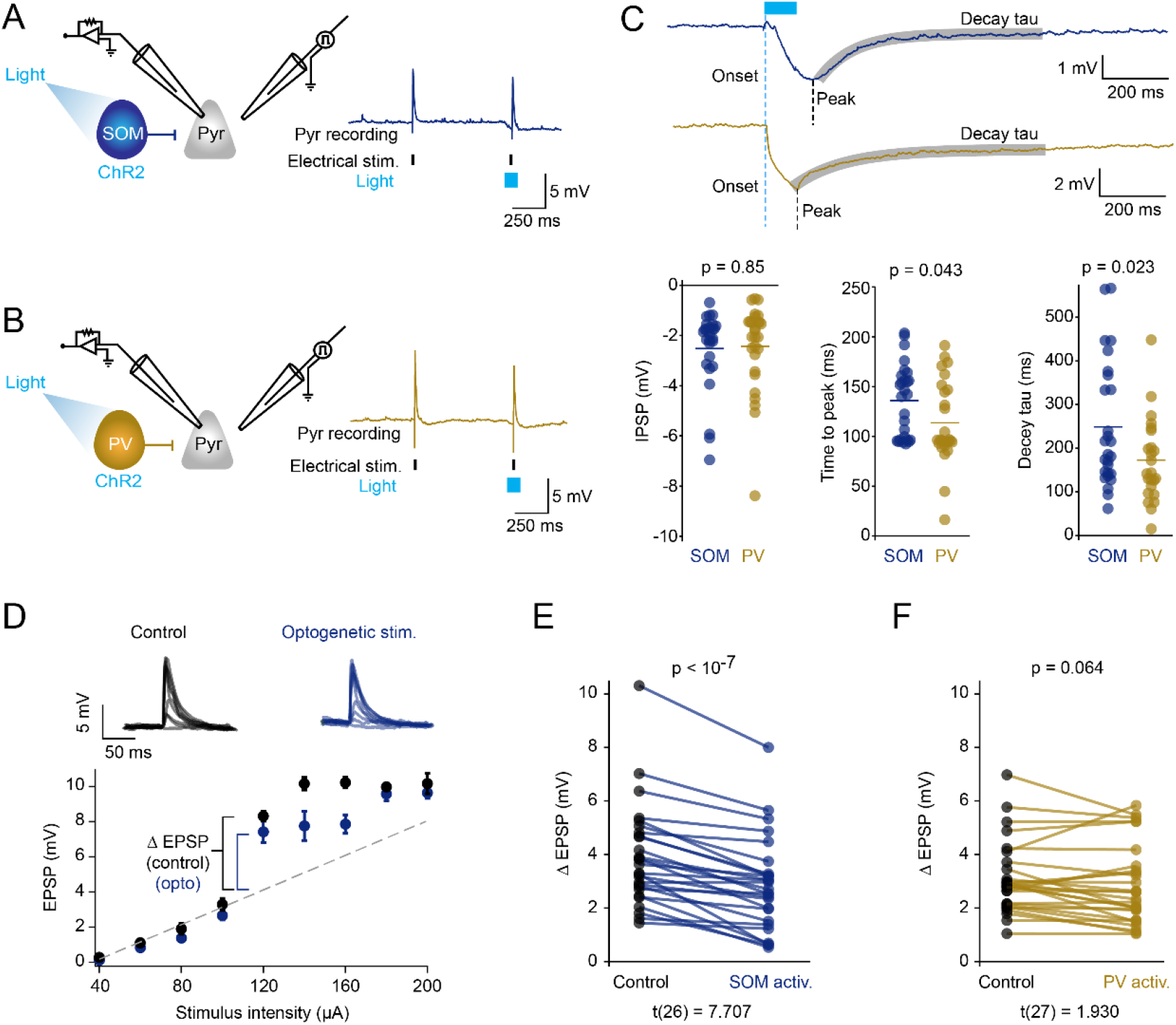
SOM activation reduces the magnitude of non-linear responses. (A) Left, cartoon schematic depicting recording and optogenetic activation of a SOM interneuron. Right, example pyramidal cell response to electrical stimulation (purple hash marks) and 100 ms optogenetic stimulation of SOM cells (light blue) (B) Same as (A) with activation of PV interneurons in gold. (n = 28). (C) Left, IPSP response in recorded pyramidal cells following optogenetic activation of either SOM (blue) or PV (gold) interneurons (n = 54), example traces above. Middle, time difference between optogenetic stimulus onset and peak (minimum) IPSP response in pyramidal cells during activation of either SOM (blue) or PV (gold) interneurons (n = 54), example traces above. Right, comparison of decay exponential (tau) in pyramidal IPSP responses from during optogenetic stimulation of either SOM (blue) or PV (gold) interneurons (n = 54). (D) Top, cartoon schematic depicting recording and optogenetic activation of a SOM interneuron subtype (blue). Middle, example voltage response to control stimulation (i.e. in the absence of optogenetic activation - black) and stimulation during optogenetic activation of SOM interneurons (blue). Bottom, example IO curve during control subthreshold stimulation (black) and during optogenetic activation of SOM interneurons (blue). Dashed line indicates linear extrapolation of first 4 mean values. Error bars indicate ± s.e.m. (E) Top, cartoon schematic depicting recording and optogenetic activation of a SOM interneurons (blue). Bottom, comparison of the magnitude of experimental EPSP response minus expected linear extrapolation under control conditions (black) to optogenetic activation of SOM interneurons (blue) (n = 27). (F) Same as (E) with PV interneuron activation in gold (n = 28)

**Figure 3.**
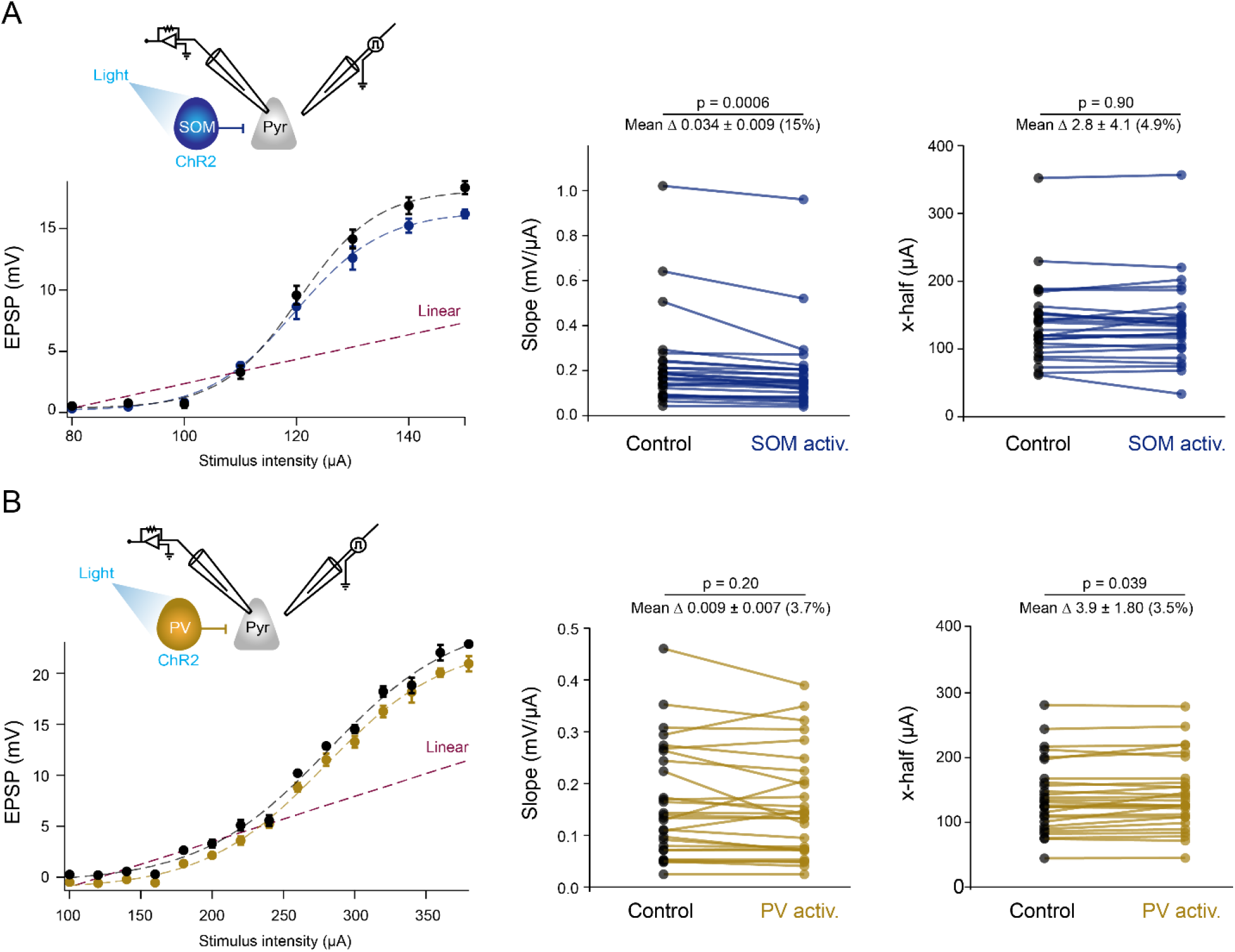
SOM cells mediate predominantly divisive gain control while PV activation results in modest subtractive inhibition. (A) Left, example IO response to linearly increasing levels of electrical stimulation, fitted with a sigmoidal curve, in the absence (black) or presence (blue) of optogenetic activation of SOM cells. Dashed line (purple) indicates linear extrapolation from first four data points. Error bars indicate ± s.e.m. Middle, comparison of slope of sigmoidal fit during control (black) to optogenetic activation of SOM cells (blue). Right, comparison of the x-half (offset) of sigmoidal fit during control (black) to optogenetic activation of SOM cells (blue). (n = 27). (B) same as (A) with activation of PV interneurons in gold (n = 28)

**Figure 4.**
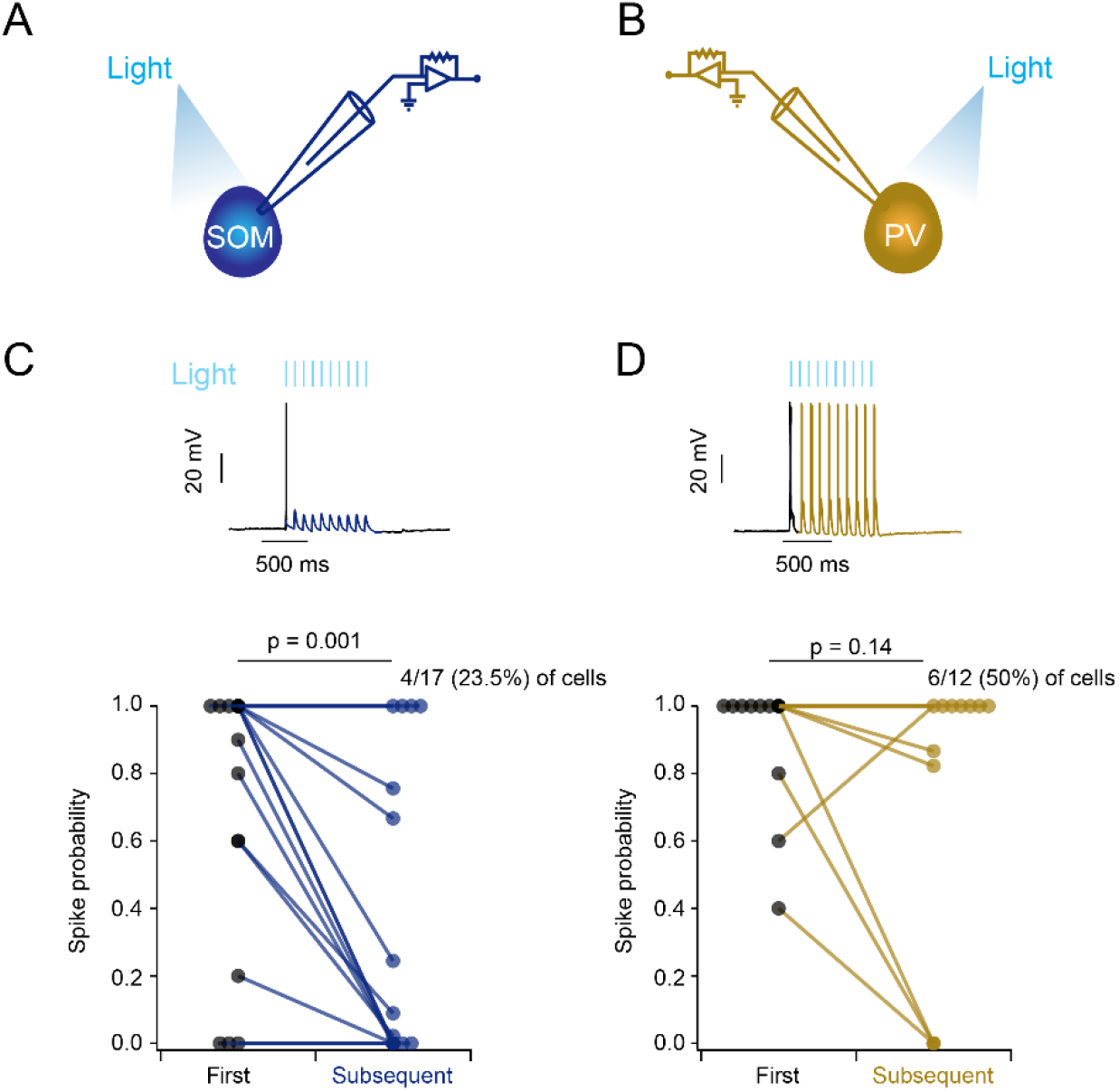
SOM cell activation reduces spike fidelity to trains of optogenetic stimuli. (A) Cartoon schematic of SOM recording configuration. (B) same as (A) with recording/activation of PV interneurons. (C) Top, example SOM cell response to 10 Hz trains of 450 nm light pulses (20 ms pulse width), initial response black, subsequent responses blue. Bottom, comparison of spike probability to the initial pulse in a 10 Hz train of light to all subsequent pulses. The majority (58.8%) of cells showed a reduction in action potential firing to subsequent pulses while only 23.5% of cells responded with equal probability to all pulses (n = 17). (D) Same as (C) but during activation of PV cells in gold. 50% of cells responded too all pulsus in a train while only 25% showed a reduced action potential probability to subsequent pulses (n = 12).

## Results

### Electrical stimulation of afferent axons results in NMDAR-dependent supralinear integration

We made whole-cell current clamp recordings from layer 2/3 pyramidal cells in slices of mouse visual cortex. To activate nonlinear mechanisms on dendrites, we electrically stimulated nearby axons with increasing stimulus intensities (**Figure 1A**). Brief (0.1 ms) current pulses from the stimulating pipette were sufficient to elicit EPSPs at the soma. The majority of pyramidal cells (55/69) exhibited a characteristic response to increasing stimulus intensity (**SI**) values, in which the EPSP values increased surpalinearly above a cell-variable SI threshold (**Figure 1B**). To identify the voltage-dependent channels contributing to the nonlinear response, we blocked NMDA receptors with the competitive antagonist (2R)-amino-5-phosphonopentanoate (**APV**) (**Figure 1C**). Bath application of APV resulted in expected reductions in the duration of the EPSP (**Supplemental Figure 5**; mean full-width at half-maximal (**FWHM**) control = 20.9 ± 2.4 ms, mean FWHM APV = 12.0 ± 1.7 ms, *p* = 0.009; mean tau control = 26.3 ± 2.9 ms, mean tau APV = 15.1 ± 2.2 ms, *p* = 0.005, n = 8). Blocking NMDA receptors resulted in a significant reduction in the slope of the input-output (**IO**) curve (**Figure 1D**), bringing the mean EPSP values closer to the linear trajectory extrapolated from the lower stimulus intensities (46.6% reduction in slope, mean slope control = 0.06 ± 0.01 mV/μA, mean slope APV = 0.03 ± 0.01 mV/μA, *p* < 0.01, n = 8). Blocking NMDA receptors did not result in a completely linear IO curve in most cells, suggesting the contribution of other active voltage-gated channels (e.g. Na^+^ or Ca^2+^ channels) to dendritic non-linearities. Thus, brief electrical excitation of axons recruits active mechanisms on the dendrites of layer 2/3 visual cortex neurons, manifesting as nonlinear increases in EPSP / SI input-output relationship.

**Figure 5.**
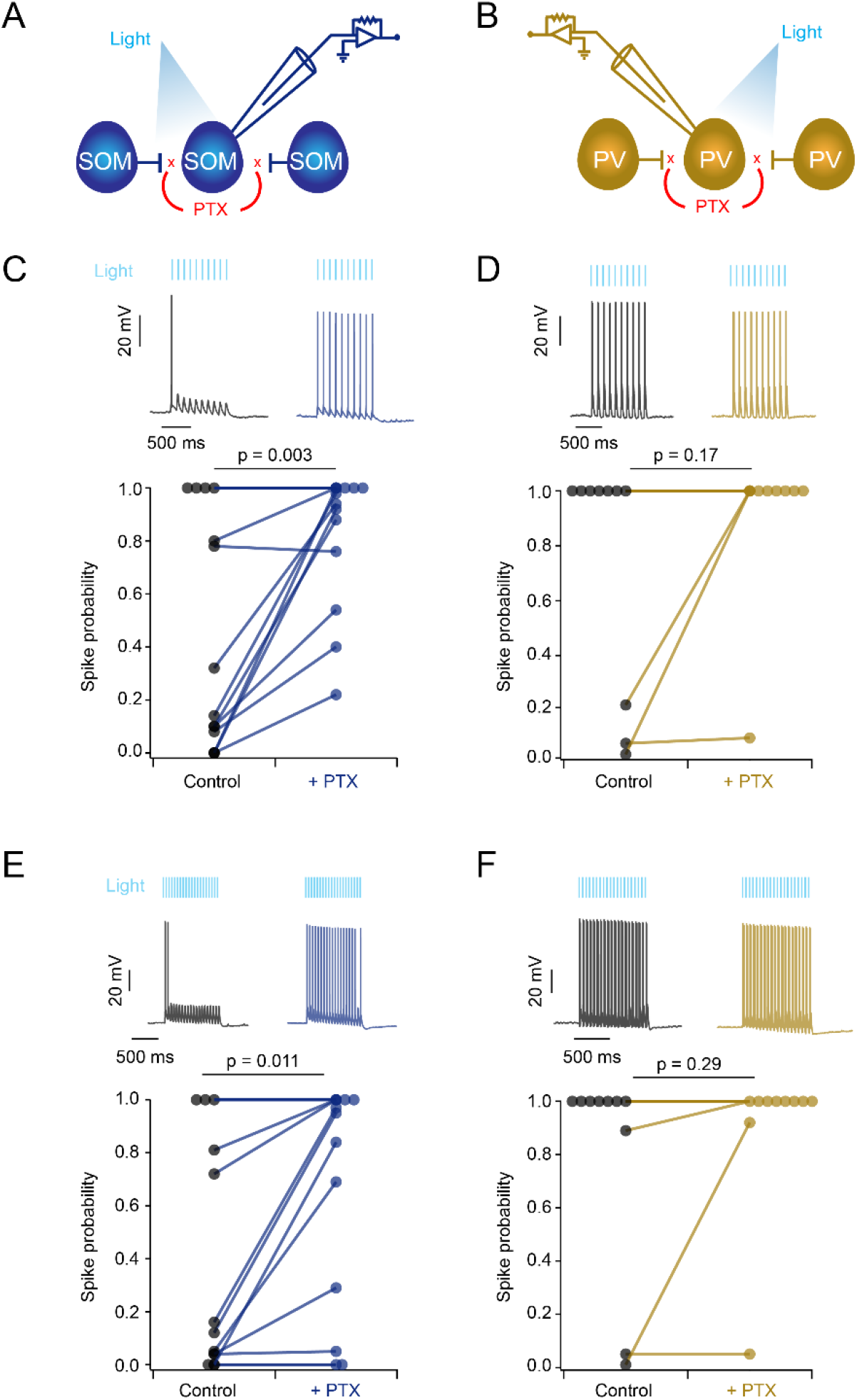
SOM cells exhibit greater GABAergic mutual inhibition than do PV cells. (A) Cartoon schematic indicating recording configuration and hypothesized mutual inhibition between SOM interneurons. (B) Same as (A) with recording/activation of PV interneurons in gold. (C) Top, example responses of SOM cell to 10 Hz trains of 450 nm light pulses (20 ms pulse width) in control aCSF (black trace) and aCSF containing 50 μM PTX (blue trace). Bottom, comparison of intracellular spike probability to all pulses in a 10 Hz train of light in control aCSF and PTX containing aCSF (n = 14). (D) Same as (C) with recording/activation of PV interneurons in gold (n = 10). (E) same as (C) showing SOM cell response to 20 Hz trains of 450 nm light pulses (n = 13). (F) same as (D) showing PV cell response to 20 Hz trains of 450 nm light pulses in gold (n = 10).

### Activation of SOM but not PV interneurons decreases EPSP amplitudes in the supralinear regime

Next, we investigated the effects of SOM and PV interneurons on active nonlinear synaptic integration. We crossed mice that expressed cre-recombinase in an interneuron subtype with animals that express an optimized form of the light-activated cation channel channelrhodopsin-2 (ChR2) and the fluorescent protein tdTomato (tdTom) in a cre-dependent fashion. Mice thus expressed both ChR2 and tdTom in either SOM (SOM-cre^+^/ChR2^+^/tdTom^+^) or PV cells (PV-cre^+^/ChR2^+^/tdTom^+^). We made whole cell recordings from pyramidal neurons, as before, in slices from either SOM or PV mice. In each sweep, electrical stimulation was presented with and without optogenetic stimulation of inhibitory neurons (SOM or PV) (**Figure 2A and B**). Optogenetically evoked inhibitory post-synaptic potentials (**IPSP**) amplitudes recorded in pyramidal neurons were similar for both SOM and PV cell activation (**Figure 2C**; SOM: mean = −2.518 ± 0.299 mV, n = 27; PV: mean = −2.432 ± 0.325 mV, n = 28; *p* = 0.846). However, they exhibited notable difference in the time course with the IPSPs resulting from stimulation of SOM cells displaying a significantly longer delay time to peak (**Figure 2C**; SOM mean = 135.9 ± 7.1 ms, n = 27; PV mean = 113.5 ± 8.1 ms, n = 28; *p* = 0.043) and slower time to return to baseline compared to those by PV cells (**Figure 2C**; SOM mean = 248.9 ± 27.7 ms, n = 27; PV mean = 172.8 ± 17.8 ms, n = 28; *p* = 0.023).

To analyze the role of each interneuron subtype in synaptic integration, we examined the degree to which activating SOM or PV interneurons affected the nonlinear IO curves. Specifically, we were interested in whether a supralinear response would be preserved during optogenetic activation and how each interneuron subtype might affect the magnitude of the response and the SI threshold value at which the IO curves became nonlinear. Most cells (55/69) exhibited a nonlinearity in their IO curve (examples of linear cells are shown in **Supplemental Figure 3**). The supralinear response was preserved during optogenetic stimulation for the majority of cells (47/55). SOM cell activation resulted in a switch from non-linear to linear IO curve in 6 of 27 cells, while PV cell activation led to linearization in just 2 of 28 non-linear cells (**Supplemental Figure 6**). Thus, SOM cells appear to exert a greater degree of influence on nonlinear synaptic integration than PV cells. However, neither interneuron subtype consistently eliminated nonlinear synaptic integration, and instead had other effects on the IO curves.

To assess the effects of SOM and PV cells on the IO curves, we first examined how optogenetic stimulation affected the magnitude of the non-linear increase. We determined the SI value at which responses became non-linear (see Methods for further details) and compared the experimental EPSP response to the expected EPSP response based on a linear extrapolation at that SI value (**Figure 2D**). We found that optogenetic activation of SOM cells resulted in a significant reduction in the magnitude of the non-linear increase compared to control (**Figure 2E**; control mean = 3.9 ± 0.4 mV, opto mean = 2.9 ± 0.3 mV, *p* = 3.9 * 10^−8^, n = 27). In contrast, activating PV interneurons during electrical stimulation did not significantly alter the magnitude of the non-linear response (**Figure 2F**, control mean = 3.4 ± 0.3 mV, opto mean = 3.0 ± 0.3 mV, *p* = 0.064, n = 28). Thus, activation of SOM, more so than PV interneurons, affect the dendritic non-linearity.

### SOM cells mediate predominantly divisive gain control while PV cells modestly affect threshold

To assess how SOM and PV cells affected the overall IO curves, we parameterized the data using sigmoidal curve fits (see Methods for further details). We found that activation of SOM interneurons strongly reduced the slope of the sigmoidal fit compared to control conditions (**Figure 3A**; mean slope control = 0.22 ± 0.04 mV/μA, mean slope opto = 0.19 ± 0.04 mV/μA, *p* = 0.0006, n = 27). Of note, it had no effect on offset, which is related to the threshold for supralinear synaptic integration. (**Figure 3A**; mean offset control = 120.0 ± 17.5 μA, mean offset opto = 118.5 ± 18.0 μA, *p* = 0.896, n = 27). By contrast, PV cell activation moderately increased the offset of the IO curves (**Figure 3B**; mean offset control = 139.0 ± 10.5 μA, mean offset opto = 142.9 ± 10.4 μA *p* < 0.039, n = 28), but had little effect on the slope (**Figure 3B**, mean slope control = 0.16 ± 0.02 mV/μA, mean slope opto = 0.16 ± 0.01 mV/μA, *p* = 0.195, n = 28). Taken as a whole, these results demonstrate that SOM interneurons have a greater effect than PV interneurons on voltage-dependent synaptic integration. SOM cells mediate a predominantly divisive form of inhibitory gain control during active synaptic integration, while PV cells appear to contribute modest subtractive inhibition during active synaptic integration.

### SOM interneurons exhibit a greater degree of mutual inhibition than PV interneurons

While determining the appropriate laser intensity and duration necessary to achieve optimal optogenetic stimulation, we found that SOM interneurons inhibit each other to a greater degree than do PV interneurons. We made whole-cell recordings directly from SOM or PV interneurons to characterize their response to optogenetic stimulation (**Figure 4A and B**). SOM and PV cells both responded to depolarizing current steps consistent with prior reports^40,41^ (**Supplemental Figure 7 and 8**). We characterized how each interneuron subtype responded to trains of light pulses (10 and 20 Hz, 20 ms pulse duration). Four SOM cells (out of 17 total) reliably responded to all pulses in a train by firing at least one action potential per pulse, while three SOM cells were non-responsive. The remaining 10 SOM cells reliably fired action potentials only to the first pulse in a train of light pulses, and responded with a lower reliability to subsequent pulses in the train (**Figure 4C**; across all cells mean action potential probability at first pulse = 0.7 ± 0.1, mean action potential probability to subsequent pulses = 0.3 ± 0.1, *p* = 0.001, n = 17). By contrast, the majority of PV cells (9 of 12 cells) faithfully responded to all pulses in a light train with at least one action potential per pulse (**Figure 4D**, mean action potential probability at first pulse = 0.9 ± 0.06, mean action potential probability to subsequent pulses = 0.7 ± 0.1, *p* = 0.143, n = 12).

To determine whether inhibitory inputs are responsible for the reduced responses to subsequent pulses in SOM neurons, we blocked gamma aminobutyric acid (**GABA**) transmission via bath application of the GABA_A_ receptor antagonist picrotoxin (**PTX**) (**Figure 5A and B**). In PTX, SOM cells were more likely to respond to all light pulses at both 10 Hz (**Figure 5C**, mean action potential probability across all pulses in standard aCSF = 0.45 ± 0.12, mean probability in PTX = 0.83 ± 0.07, *p* = 0.003, n = 14) and 20 Hz (**Figure 5E**, mean probability standard aCSF = 0.38 ± 0.12, mean probability in PTX = 0.68 ± 0.12, *p* < 0.011, n=13). The reduced inhibition present in PTX containing aCSF was even sufficient to induce firing in some cells that had been entirely non-responsive to optogenetic stimulation in standard aCSF. Thus, it appears that optogenetic stimulation of SOM cells leads to direct release of GABA from nearby SOM cell axons (even in the absence of somatic action potentials), sufficient to inhibit responses to light pulses, an effect which is alleviated with PTX. On the other hand, PTX had little to no effect on spike probability of PV interneurons at either 10 Hz (**Figure 5D**, mean action potential probability in standard aCSF = 0.74 ± 0.14, mean in PTX = 0.91 ± 0.09, *p* = 0.165 n = 10) or 20 Hz (**Figure 5F**, mean probability in standard aCSF = 0.80 ± 0.13, mean probability in PTX = .90 ± 0.09, *p* = 0.289, n = 10). Thus, SOM interneurons exhibited stronger mutual inhibition than PV interneurons.

## Discussion

Optogenetic manipulation of the two largest interneuron subtypes revealed their distinct inhibitory effects on nonlinear synaptic integration. Activating SOM cells reduced the magnitude of the somatic depolarization during nonlinear synaptic integration (**Figure 2**). In contrast, activating PV cells had only modest effects on nonlinear integration (**Figure 3**). These results generally support the hypothesis that SOM cells, with their more dendrite-biased axonal projection patterns, have a stronger influence over nonlinear integration than PV cells. However, SOM cell activation did not shift the threshold level of synaptic input required to activate nonlinear synaptic integration. Thus, SOM cells do not regulate the recruitment of nonlinear integration mechanisms, although they do modulate the amplitude of the resulting postsynaptic depolarization seen in the soma, placing them downstream of the dendritic mechanism for the nonlinear enhancement of the synaptic inputs, namely dendritic spikes^6^.

### Location dependent effects of inhibition

The location of inhibition relative to excitation plays a critical role in synaptic integration^23^. In the absence of active dendritic mechanisms, inhibition is most effective at modulating excitatory conductances when inhibitory sources are positioned proximal to the site of excitation^23–26^. In the presence of active dendritic mechanisms, the location-dependence of inhibition is still strong. It has been previously shown that focal GABA iontophoresis targeting perisomatic areas during nonlinear responses to glutamate uncaging^29^ results in a reduction in the overall magnitude of supralinear responses, while dendritically located GABA iontophoresis leads to a shift in the stimulus intensity threshold for supralinear responses^29^. Once the increased threshold is reached, however, the magnitude of the somatic depolarizations remains comparable^29^. These reports support the notion of distinct computational roles for proximal and distal inhibition.

Although SOM and PV interneurons have relatively distinct projection patterns, the differences are more subtle than the contrast that can be achieved with local GABA iontophoresis, as used in prior work^29^. Even at dendritic locations, PV inputs can outnumber SOM inputs^42^. However, our results show that, on average, SOM inputs were distal relative to PV inputs. Peak IPSP responses resulting from stimulation of SOM cells were significantly later in arriving to the soma than IPSPs evoked from stimulating PV cells (**Figure 2C**), implying a more distal origin. Despite this, our results differed markedly from predictions from modeling and iontophoresis studies, and this prompts a re-evaluation of physiological roles for SOM and PV particularly in the context of nonlinear synaptic integration.

We found that SOM-mediated inhibition functions both as a restrictor on the absolute charge conveyed to the soma and as a gain modulator, altering the slope of the IO curve. The divisive effect of activating SOM cells reduced not only the slope of the IO plots, but also the magnitude of active dendrite dependent EPSPs (the first nonlinearity step; **Figure 2E**) measured at the soma. The inhibition mediated by PV cells had only modest effects on the offset (i.e., threshold) of the IO curves (**Figure 3B**). Thus, our findings are more in line with a similar recent experiment in hippocampal pyramidal cells from area CA1, in which a combination of two-photon glutamate uncaging and one-photon GABA uncaging demonstrated that dendritic inhibition was more effective than somatic inhibition at shunting non-linear dendritic responses^31^. Similarly, we found that inhibition mediated by SOM cells (putatively distal relative to PV interneurons) suppresses supralinear responses in the soma, without affecting the input threshold for the generation of supralinear responses. This may be a result of differences in the IPSP generated by each interneuron subtype. SOM cell IPSPs took longer to reach peak magnitude due to the distal location of their projections, however the IPSP response at the soma was similar in magnitude for both SOM and PV activation (**Figure 2C**). Due to attenuation of charge during propagation, this implies that the IPSP response experienced at the dendrites was likely greater during SOM stimulation than PV stimulation. If the majority of SOM inputs were both larger and more distal than PV inputs, yet proximal relative to excitatory sources, this could explain how the non-linear response was reduced in magnitude and gain in response to SOM but not PV stimulation, and account for the lack of effect on the initiation of dendritic non-linearities.

There are two main limitations to our study. The first concerns the method used to induce dendritic non-linearities. The use of electrical stimulation of presynaptic axons to recruit nonlinear synaptic integration is realistic in that it uses physiological synapses (rather than glutamate uncaging), but it also limits our ability to spatially control the location of synaptic excitation relative to inhibition. We targeted electrical stimulation to distal regions of the recorded cell’s dendritic arbor (**Figure 1B**), the site for dendritic spike generation in layer 2/3 pyramidal cells of the visual cortex^6^. However, it is possible that *in vivo* patterns of excitation and inhibition have a precise architecture^43^ that our experiments failed to capture. It is also possible that SOM axons were activated in addition to excitatory axons during electrical stimulation, altering the baseline SOM activity prior to optogenetic stimulation. The second limitation is with the optogenetic stimulation which is relatively strong (few if any SOM neurons failed to respond to the light) and may be considered an upper limit case. Note that the effects on nonlinear synaptic integration were modest even with this strong activation of inhibitory inputs. Thus more modest stimulation regimes would be expected to yield smaller changes than reported here. Overall, these limitations are important to consider, but leave the qualitative results we report intact.

In summary, we find that the roles of SOM and PV inhibition do not map neatly onto the roles suggested by prior work for dendritic and somatic-targeted inhibition. Our results demonstrate that SOM-mediated inhibition reduces the amplitude of somatic EPSPs during supralinear synaptic integration, and PV-mediated inhibition does not. More importantly, neither SOM nor PV inhibition cause large changes in the threshold for the recruitment of nonlinear mechanisms.

### Mutual inhibition among SOM interneurons

Our experiments also revealed strong mutual inhibition among SOM cells, and relatively weak inhibition among PV cells (**Figure 5**). These results are surprising for two reasons: recent studies on mutual inhibition between interneuron subtypes have shown that SOM cells inhibit most other interneuron subtypes while avoiding inhibiting one another (but see ref^44^); and PV cells exhibit the opposite connectivity patterns, strongly inhibiting other PV interneurons while making few connections to other genetically distinct interneurons^45^.

Several possibilities could account for the discrepancy between our results and the presumed lack of connectivity between SOM cells from prior studies. A previous connectivity study examined IPSCs in genetically distinct interneuron subtypes to a single short pulse of optogenetic activating light^45^. Trains of pulses might have revealed the connectivity we observed. Also, SOM cells are not a uniform class^46^. Different sampling biases could partially explain the discrepancy. In this work, immunohistochemistry confirmed the presence of somatostatin in the vast majority of the population of tdTom^+^ cells (**Supplemental Figure 9**). With these points, a parsimonious explanation for our results is that the broad population of SOM cells exhibit more mutual inhibition than previously thought.

In summary, our results provide new evidence for mutual inhibition among SOM interneurons. Inhibition from these SOM interneurons suppresses nonlinear synaptic integration more than that from PV neurons. However, neither SOM nor PV interneurons significantly shifts the threshold for nonlinear dendritic integration.

---

**Supplemental Figure 1.**
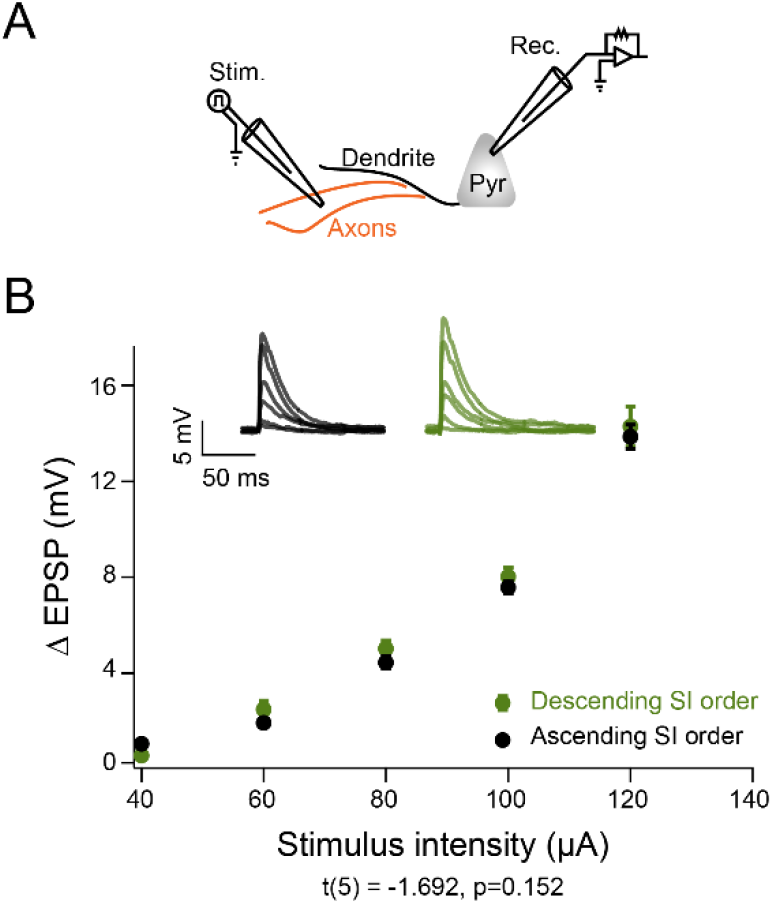
Stimulus intensity presentation does not influence EPSP magnitude. (A) Cartoon schematic of recording configuration (B) Example EPSP response to linearly increasing (black) or decreasing (green) stimulus intensities. Inset, sample voltage traces.

**Supplemental Figure 2.**
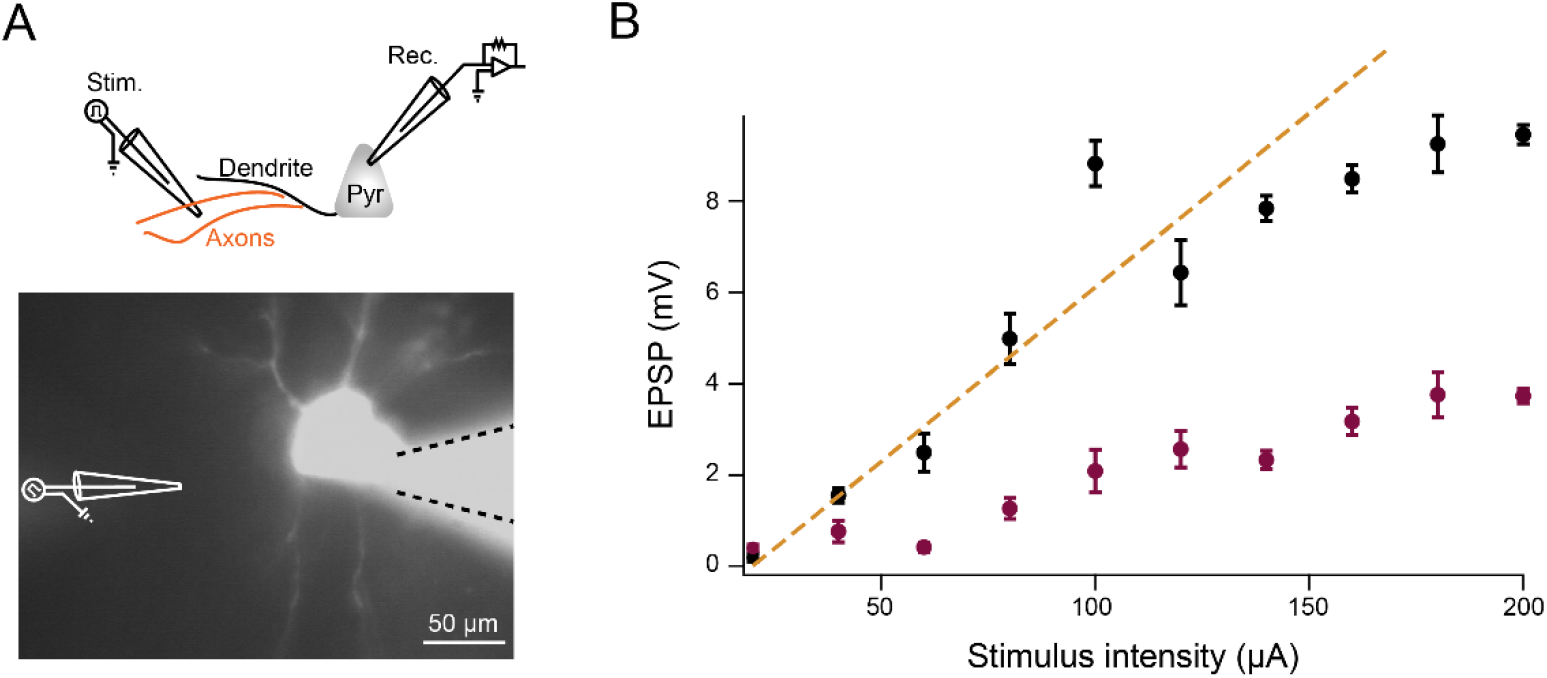
Sublinear pyramidal cell. (A) Top, cartoon schematic of recording configuration. Bottom, example infrared image of layer 2/3 pyramidal cell filled with fluorescent Alexa 594 dye. (B) Input-out plot showing sublinear response to linearly increasing current stimulations in control aCSF (black) or aCSF containing 100 μM APV (purple)

**Supplemental Figure 3.**
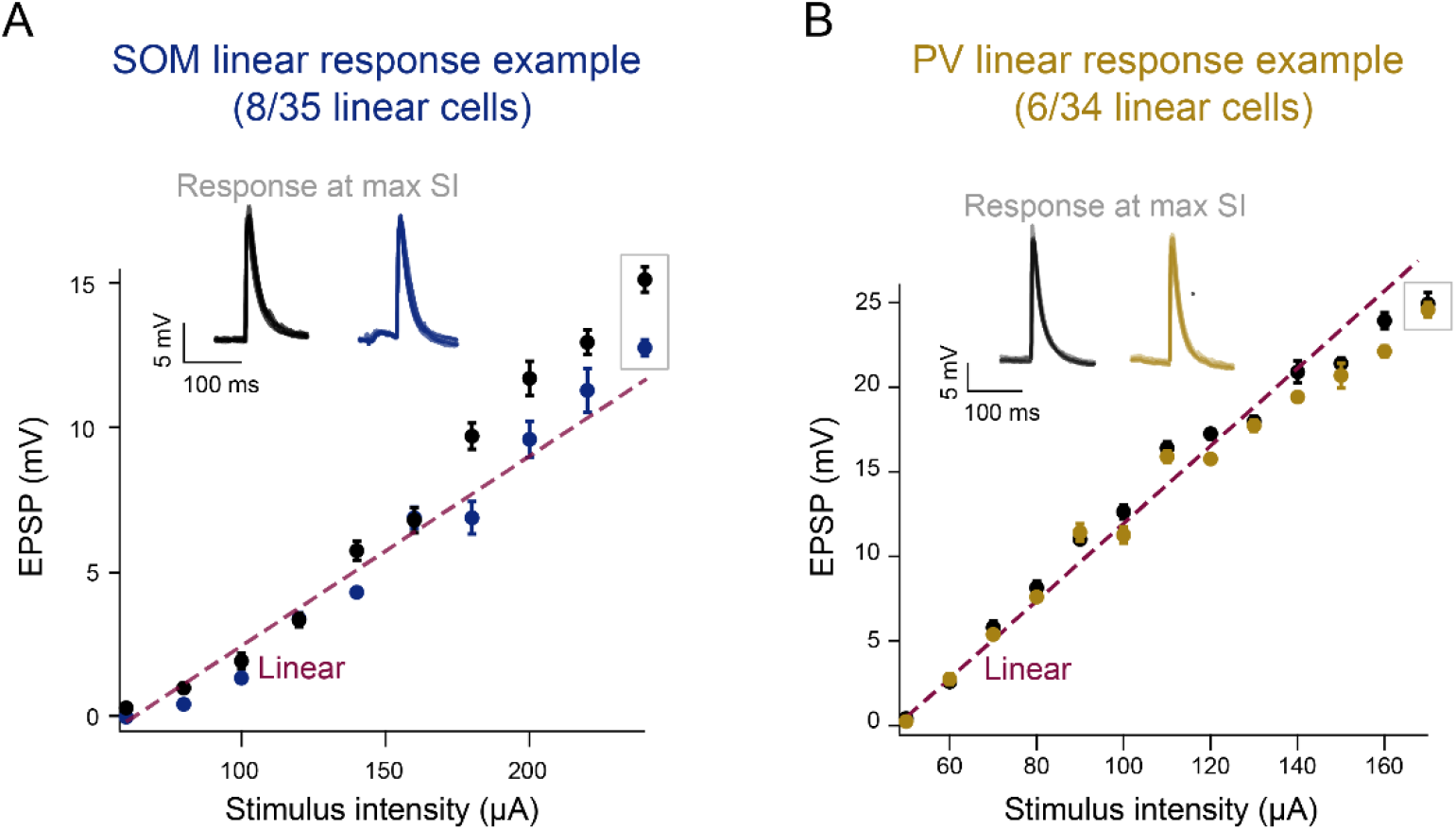
Example linear responding cells. (A), example linear IO response to increasing levels of electrical stimulation in the absence (black) or presence (blue) of optogenetic activation of SOM cells. Inset, sample voltage traces at maximal stimulus intensity. Dashed line (purple) indicates linear extrapolation. Error bars indicate ± s.e.m. (B), same as (A) with PV activation in gold.

**Supplemental Figure 4.**
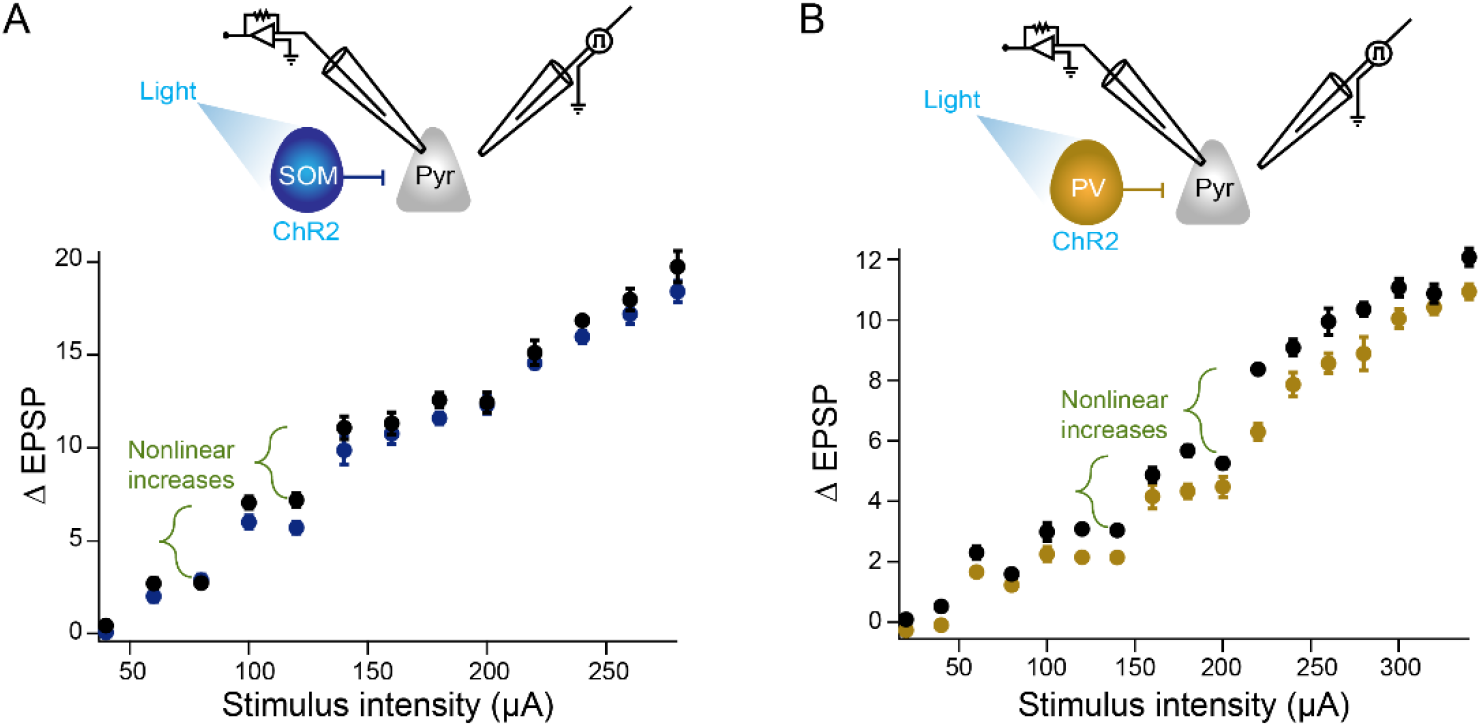
Example cells with multiple non-linear increases. (A) Top, cartoon schematic depicting recording and optogenetic activation of a SOM interneuron. Bottom, example input out trace during control stimulations (black) or stimulations during optogenetic activation of SOM interneurons (blue). (B) Same as (A) with activation of PV interneurons in gold.

**Supplemental Figure 5.**
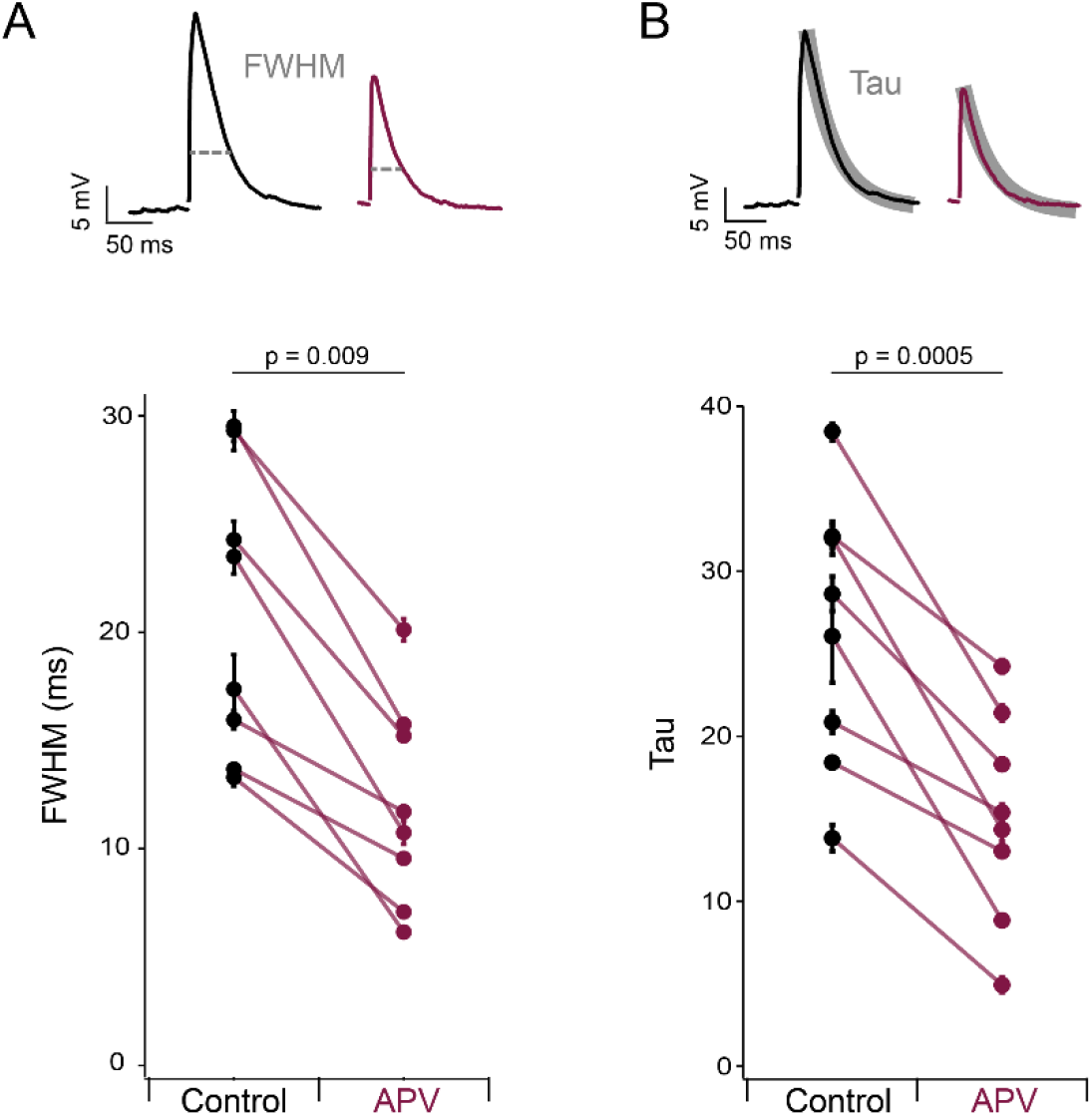
APV affect EPSP kinetics. (A) Bottom, comparison of the full width at half max (FWHM) of induced EPSP at maximal stimulus intensities in control aCSF (black) or aCSF containing 100 μM APV (purple). Top, example voltage traces. (B) Same as in (A) with the comparison of the decay constant, tau, in control (black) and APV containing (purple) aCSF. (n = 8).

**Supplemental Figure 6.**
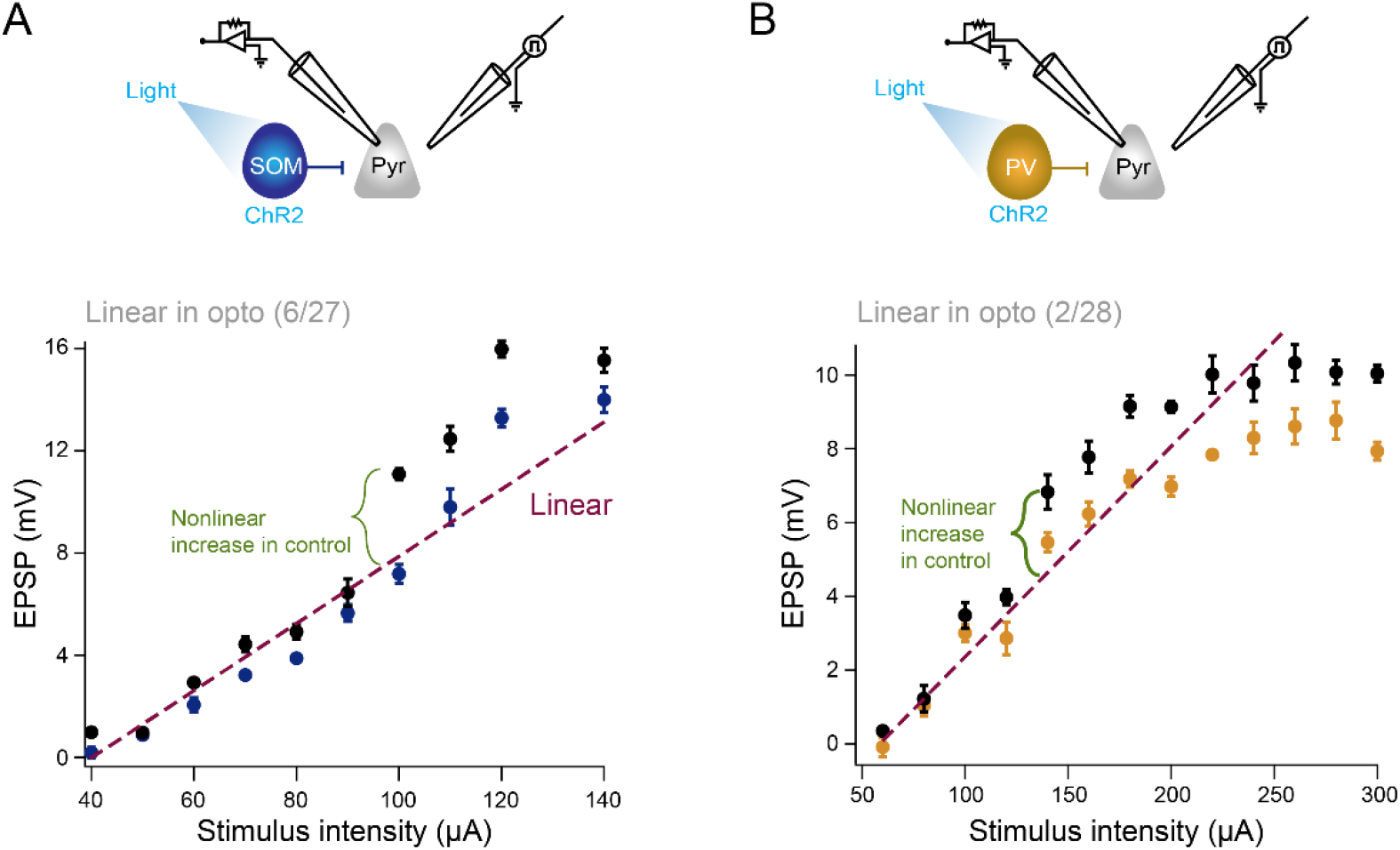
Example cells with non-linear increases during control stimulation that were linearized during optogenetic stimulation. (A) Top, cartoon schematic depicting recording and optogenetic activation of a SOM interneuron. Bottom, example input out trace during control stimulations (black) or stimulations during optogenetic activation of SOM interneurons (blue) Dashed line (purple) indicates linear extrapolation. (B) Same as (A) with activation of PV interneurons in gold.

**Supplemental Figure 7.**
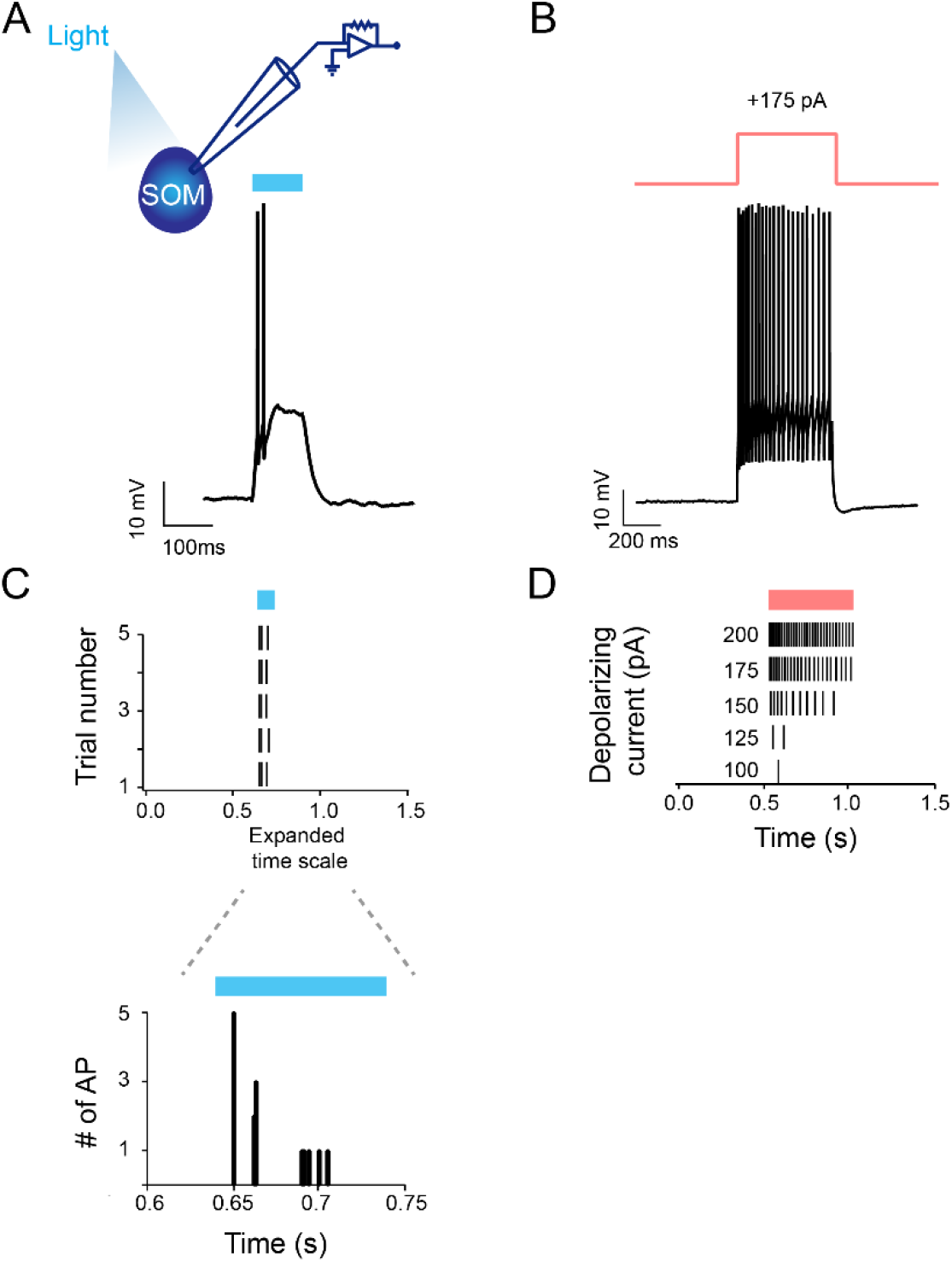
SOM cells respond with high rate of action potential firing when nearby SOM cells are not stimulated. (A) Top, cartoon schematic indicating recording configuration, Bottom, example response to 100 ms of 450 nm light pulse. (B) same cell’s response to 500 ms of +175 pA current. (C) Top, spike time raster for same cell in response to 100 ms light pulse. Bottom, peri-stimulus time histogram. (D) Spike time raster of cells response to increasing steps of depolarizing current.

**Supplemental Figure 8.**
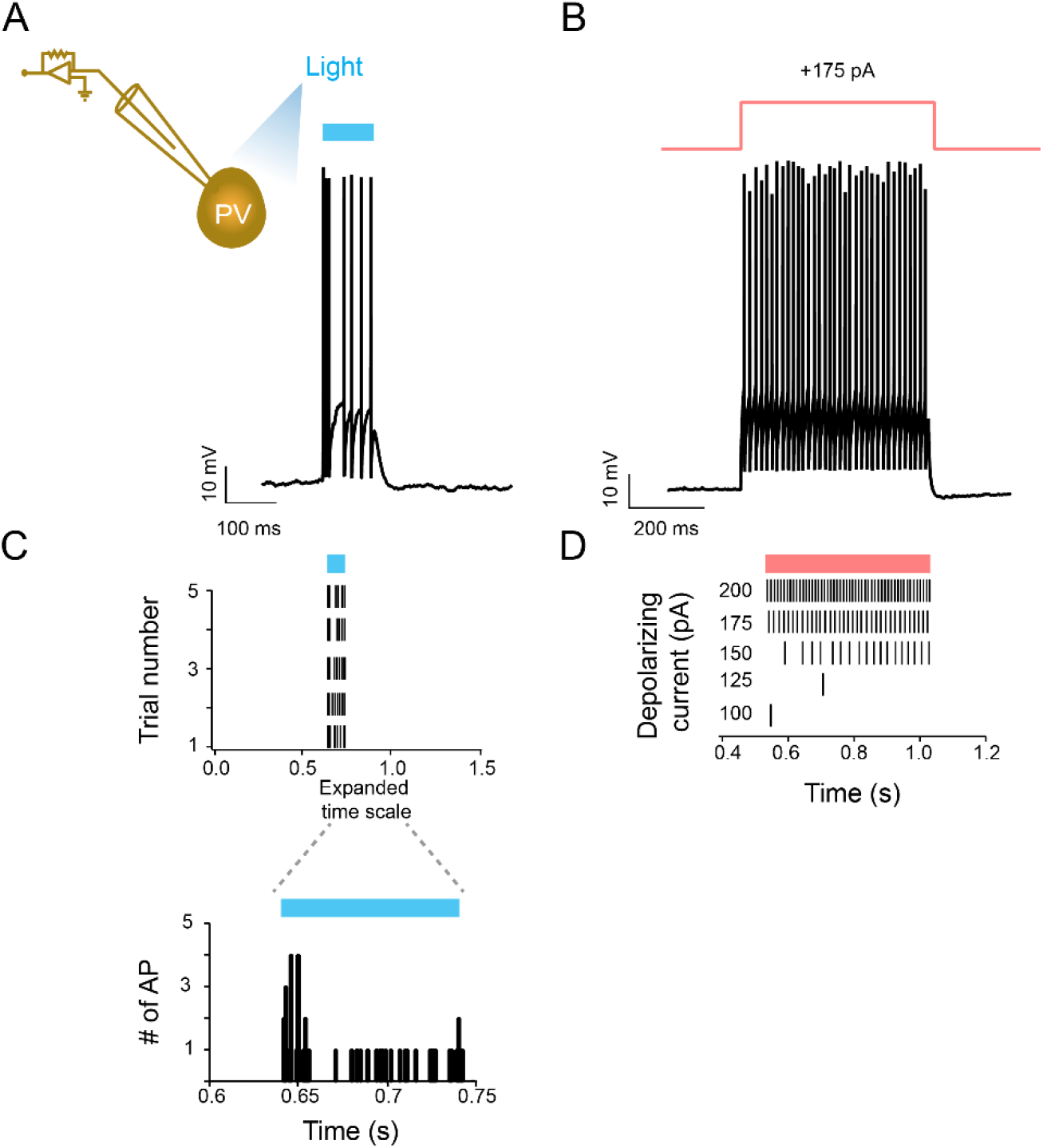
PV cells respond with high rate of action potential firing when nearby PV cells are not stimulated. (A) Top, cartoon schematic indicating recording configuration, Bottom, example response to 100 ms of 450 nm light pulse. (B) same cell’s response to 500 ms of +175 pA current. (C) Top, spike time raster for same cell in response to 100 ms light pulse. Bottom, peri-stimulus time histogram. (D) Spike time raster of cells response to increasing steps of depolarizing current.

**Supplemental Figure 9.**
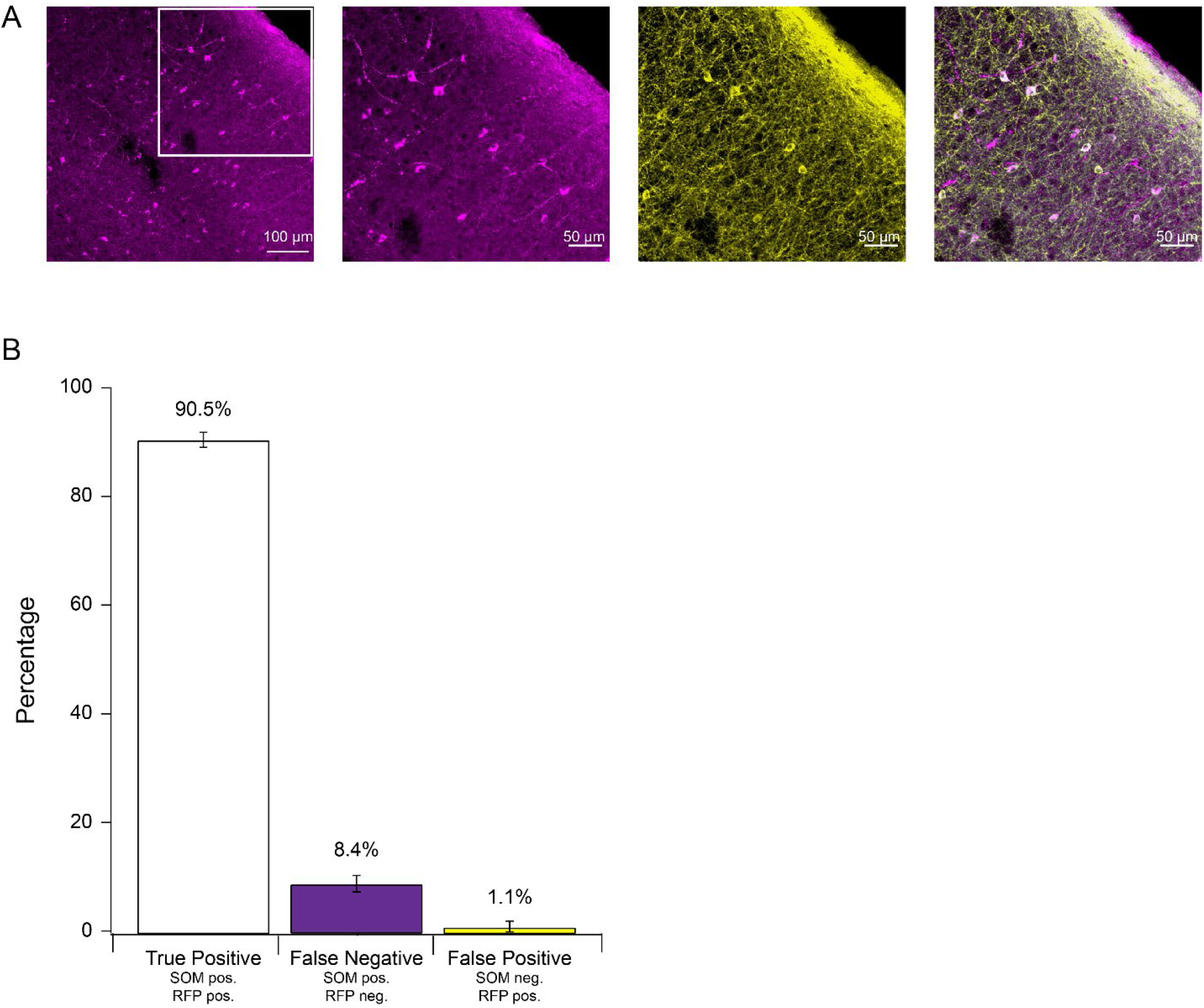
The majority of tdTom^+^/ChR2^+^ cells are SOM^+^. (A) Left, representative image of coronal section of mouse visual cortex stained with somatostatin and tdTomato antibodies. Middle-left, enlarged section of leftmost image showing cells positive for somatostatin expression. Middle-right, same as middle-left showing cells positive for tdTomato expression. Right, merged image. (B) Quantification of cell expression, mean ± s.e.m. (6 sections, 2 mice)

